# Contralateral delay activity and alpha lateralization reflect retinotopic and screen-centered reference frames in visual memory

**DOI:** 10.1101/2023.06.19.545421

**Authors:** Wanja A. Mössing, Svea C. Y. Schroeder, Anna Lena Biel, Niko A. Busch

## Abstract

The visual system represents objects in a lateralized manner, with contralateral cortical hemispheres responsible for left and right visual hemifields. This organization extends to visual short-term memory (VSTM), as evidenced by electrophysiological indices of VSTM maintenance: contralateral delay activity (CDA) and alpha-band lateralization. However, it remains unclear if VSTM represents object locations in gaze-centered (retinotopic) or screen-centered (spatiotopic) coordinates, especially after eye movements. In two experiments, participants encoded the colors of target objects and made a lateral saccade during the maintenance interval, thereby shifting the object’s location on the retina. A non-lateralized probe stimulus was then presented at the new fixation for a change detection task. The CDA maintained lateralization towards the target’s original retinotopic location, unaffected by subsequent saccades, and did not invert polarity even when a saccade brought that location into the opposite hemifield. We also found conventional alpha lateralization towards the target’s location before a saccade. After a saccade, however, alpha was lateralized towards the screen center regardless of the target’s original location, even in a control condition without any memory requirements. This suggest that post-saccadic alpha-band lateralization reflects attentional processes unrelated to memory, while pre- and post-saccade CDA reflect VSTM maintenance in a retinotopic reference frame.

## 1 Introduction

The visual system is fundamentally organized in a lateralized fashion, whereby objects located in the left or right visual hemifield are processed in the contralateral cortical hemisphere (Wandell, Dumoulin, & Brewer, 2007). This organization applies not only to perception, but also to cognitive processes operating on visual information. Specifically, covert spatial attention towards a lateral location modulates activity primarily in contralateral visual cortex (Heinze et al., 1994; Tootell et al., 1998). Likewise, visual short-term memory (VSTM) representations of objects that are no longer in view are stored in the hemisphere contralateral to the object’s original location (Brincat et al., 2021). The two hemispheric VSTM stores even seem to operate largely independently, as the number of memorized objects in one hemifield has little effect on storage capacity (Delvenne, 2005; Umemoto, Drew, Ester, & Awh, 2010) and neural correlates of memory load (Kornblith, Buschman, & Miller, 2016) in the opposite hemisphere. However, the fact that humans make eye movements as often as three times per second (Ibbotson & Krekelberg, 2011), and that each eye movement changes the location or even the hemifield of the memorized objects, begs the question of the nature of “location” in VSTM: after an eye movement, are memories still bound to the hemisphere contralateral to the object’s original location on the retina, or are they updated according to its current spatiotopic location on the screen? Intuitively, our experience of visual stability across eye movements suggests that visual information is at some point transformed into a spatiotopic representation. However, Golomb and Kanwisher (2012b) have demonstrated that subjects were better at reporting a memorized target’s pre-saccadic retinotopic location than its updated spatiotopic location, and similar results have been found in peceptual and attentional tasks (Golomb, Chun, & Mazer, 2008; Golomb & Kanwisher, 2012a). Does this predominance of a retinotopic reference frame in VSTM also apply to lateralized electrophysiological markers of VSTM maintenance?

The contralateral delay activity (CDA) is a sustained negative event-related potential that occurs during the maintenance interval at parietal-occipital electrodes contralateral to the memorized items. Its amplitude increases with the number of memorized objects, reaching an asymptote at participants’ individual VSTM capacity (Vogel & Machizawa, 2004; Vogel, McCollough, & Machizawa, 2005). Importantly, this signal carries information not only about the objects’ original spatial location, but also about their non-spatial properties such as orientation (Bae & Luck, 2018). Hence, the CDA is regarded as a neural signature of active storage in VSTM, closely linked to the actual memory content (Luria, Balaban, Awh, & Vogel, 2016).

Furthermore, the power of neural oscillations in the alpha-band (approximately 8 to 12 Hz) decreases during the maintenance interval at posterior electrodes contralateral to the memorized items (Sauseng et al., 2009). Thus, the pattern of alpha-band lateralization during VSTM maintenance appears similar to attention-induced alpha lateralization, where alpha power decreases contralateral to the attended hemifield (Kelly, Lalor, Reilly, & Foxe, 2006; Thut, 2006). While alpha lateralization carries information about the memoranda’s locations (Foster, Sutterer, Serences, Vogel, & Awh, 2016) similar to the CDA, it cannot be used to decode non-spatial object properties (Bae & Luck, 2018). Hence, alpha lateralization during VSTM maintenance is often characterized as a support process, i.e. sustained spatial attention towards a memory representation (van Ede, 2018).

CDA and alpha lateralization are typically studied in change detection tasks, in which subjects memorize features (e.g. color) of lateralized target objects, maintain this information during a short delay, and then judge whether a probe stimulus is identical to the memorized object. Critically, subjects are usually instructed to maintain central fixation throughout the entire trial, precluding a dissociation between retinotopic and spatiotopic reference frames. Here, we employed an adaptation of this paradigm, in which subjects encoded lateralized target objects and memorized them during a delay interval. Halfway through the delay interval, they were cued to make a saccade to the left or the right, or to maintain fixation. This allowed us to test whether CDA and alpha lateralization after the saccade reflects the memoranda’s original retinotopic locations, or their updated spatiotopic location.

## 2 Materials and Methods

### 2.1 Participants

Thirty subjects were included in the analysis of experiment 1 (17 female; aged 19–35 years, 27 right-handed). Data from another two subjects were not included because their eye tracking data were corrupt. Thirty-five subjects were included in the analysis of experiment 2 (27 female; aged 18–27 years, 31 right-handed). Data from another five subjects who did not complete the memory capacity test were not included in the analysis. The sample sizes were not based on a systematic power analysis. Given that our main research question concerned the mere presence of a post-saccadic lateralized EEG response (CDA) and its polarity, rather than differences in CDA amplitude between experimental conditions, these sample sizes are in accordance with Ngiam, Adam, Quirk, Vogel, and Awh’s (Ngiam et al., 2021) recommendation of a minimum of 25 subjects.

All subjects had normal or corrected-to-normal vision. All subjects provided written informed consent and were compensated for participation with course credit or money (10 €/h). The study was approved by the ethics committee of the faculty of psychology and sports sciences at the University of Münster (#2018-37-WM).

### 2.2 Apparatus

Stimuli were generated using Matlab 2018b (www.mathworks.com) and the Psychophysics Toolbox 3 (Brainard, 1997; Kleiner et al., 2007; Pelli, 1997), and displayed on a 24” Viewpixx/EEG LCD Monitor with 120 Hz refresh rate, 1 ms pixel response time, 95% luminance uniformity, and 1920*×*1080 pixels resolution (www.vpixx.com). Participants were seated in a dimly-lit, sound-attenuated cabin with their head rested on a chin rest. Distance between participants’ eyes and monitor was approximately 86 cm. Participants responded using a wired Logitech F310 Gamepad.

### 2.3 Stimuli

#### Experiment 1

Stimuli were presented on a uniform gray background. Three black diamond-shaped markers (0.6° diagonals) located at the screen center and at 12° eccentricity to the left and right were present on screen throughout the experiment (Figure 1A).

**Figure 1:**
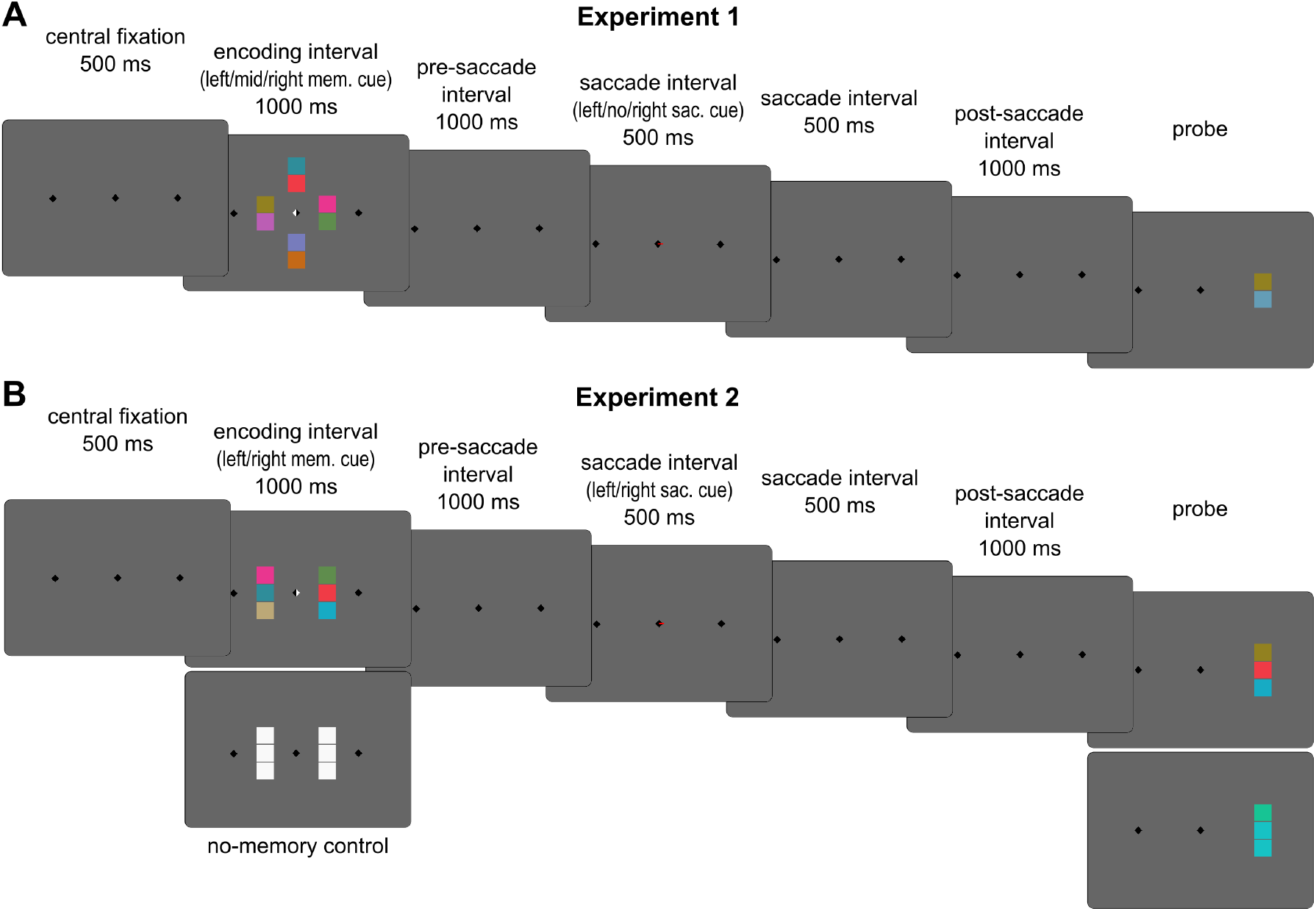
Schematic trial sequence. **A**: experiment 1; subjects initiated a trial by maintaining central fixation for 500 ms. The encoding display comprised four target objects, each consisting of two colored squares, one of which was cued for memorization (indicated by white half of diamond). After the pre-saccade delay interval, a saccade cue (thin red line) instructed subjects to make a speeded saccade to the left or the right fixation marker within 1000 ms, or to maintain central fixation. After the post-saccade interval, a probe object appeared at the new fixation and subjects localized the square whose color had changed. **B**: experiment 2; procedures were similar to experiment 1, except that subjects memorized either three colors or no color at all in the no-memory control condition. The probe in the control condition consisted of two squares with identical color and one square of slightly different color, and subjects indicated the square with the deviant color.

The target display comprised four objects at 3° eccentricity to the top, left, right, and bottom of the central diamond. Each object consisted of two vertically aligned colored squares (1° side lengths) separated by a 0.1° wide gap. Colors were chosen from a circle (radius 60°) cut out of the CIE Lab color space, centered at L = 70, a = 20, and b = 38 (Zhang & Luck, 2008). For each of the four objects, one square was colored with a value selected at random from the color circle with the restriction that colors differ by at least 30° across objects. The other square of each object was colored with the value 180° across the color circle.

After the delay intervals, a probe object consisting of two colored squares with the same size as the target objects appeared at the new fixation location, temporarily replacing the fixation marker at that location. One of the two squares had the same color as the corresponding square of the cued target object. The color of the other square was determined by a staircase procedure (Watson & Pelli, 1983), which adjusted change detection difficulty by setting the color difference between target square and probe square to a value yielding 82% detection accuracy.

#### Experiment 2

Stimuli were identical to experiment 1 except for the following changes. Only the objects to the left and right of the central diamond were presented; no objects were shown on the midline. Each object consisted of three colored squares, except for the new “no-memory” control condition, in which all squares of the target display were white. The probe stimulus in the no-memory condition consisted of three colored squares, two of which had the identical color selected at random from the color circle. The third square had a slightly different color, and the color difference was adjusted by a staircase procedure such that the deviant color could be localized with 90% accuracy (Figure 1B).

### 2.4 Procedure

#### 2.4.1 Experiment 1

Subjects initiated a trial by fixating the central diamond. After 500 ms of stable fixation, the encoding display appeared for 1000 ms. At the same time, either the left, right, top, or bottom half of the central marker turned white, instructing subjects to memorize all colors of the cued object while maintaining fixation on the central marker. Following the offset of the encoding display, subjects kept fixating the central marker throughout the 1000 ms pre-saccade delay interval. This was followed by a 1000 ms saccade-interval. A saccade cue (red line of 0.036° width) appeared during the first 500 ms of this interval. On some trials, the red line ran across the full length of the fixation marker, instructing subjects to keep fixating on the central marker. On other trials, the red line pointed to the left or right, instructing subjects to make a saccade to the fixation marker on the cued side. Subjects were required to re-fixate on the cued marker before the end of the saccade-interval, i.e. within 1000 ms after saccade-cue onset. This was followed by the final 1000 ms post-saccade delay interval, during which subjects were required to keep steady fixation at the new location. In sum, the total delay interval between offset of the encoding display and probe onset was 3000 ms. After the delay intervals, a probe object appeared at the new fixation and subjects performed a change localization task by indicating the location of the color change using the up and down buttons on the gamepad’s left side or the Y and A buttons on the right side (counterbalanced across participants). Following the button press, the selected square was surrounded for 300 ms by a green or red frame for correct and incorrect responses, respectively. After incorrect responses, the correct choice was additionally surrounded by a white rectangle. This feedback was followed by a 750 ms inter-trial interval. Trials with faulty fixation or with incorrect or tardy saccades were aborted and repeated.

Prior to recording, participants completed 15 training trials with feedback. The main experiment included 192 trials each for left and right memory-cues and 48 trials each for top and bottom memory-cues (not counting aborted trials). Each saccade cue condition (left, no saccade, right) was used on one third of trials per memory-cue condition (in pseudo-random order). A complete session lasted for approximately 150 minutes, including EEG preparation.

#### 2.4.2 Experiment 2

The procedure was identical to experiment 1 except for the following changes. The no-saccade condition was not included. Furthermore, the condition that required memorization of a target on the central midline was dropped. Instead, a new “no-memory” control condition was introduced, in which the target objects were all white and the central diamond remained black during the target interval, instructing subjects not to memorize anything. The experiment comprised 80 trials for each combination of memory-cue and saccade-cue in pseudo-randomized order. In this condition, the probe consisted of two identically colored squares and one square of a slightly different color. Subjects were instructed to indicate the location of the deviant color.

Individual short-term memory capacity was assessed in a separate block of trials after the main experiment, following procedures described in Brady, Störmer, and Alvarez, 2016 and Quirk, Adam, and Vogel, 2020. Since estimates of memory capacity are bound between zero and the set size tested, they are only valid if the set size is at least as large as an individual’s true capacity (Rouder, Morey, Morey, & Cowan, 2011). However, using too large a set size might be demotivating for subjects with substantially lower capacity. Thus, on separate trials we used either a larger (7 items) or moderate (4 items) set size, randomized across trials. On each trial, a target display was presented for 1000 ms, comprising four or seven colored discs (2° diameter), equally spaced on an imaginary circle of 4° diameter around a central fixation marker on a grey background. The discs’ colors were sampled at random, but evenly spaced around the color circle (see above) to make them as distinctive as possible. After a 800 ms delay interval, a probe disc (2° diameter) was presented at the location of one of the target discs. The upper or lower half (counterbalanced across trials) of the probe disc had the same color as the target disc at this location, while the other half was colored exactly 180° opposite in color space. Subjects indicated whether the upper or lower half-disc represented the target’s original color. No eye tracking or EEG data were recorded during the capacity test. Memory capacity *K* was computed separately for each of the two set sizes *N* as *K* = *N* (2*p* − 1), where *p* is the proportion of correct responses (Brady et al., 2016; Quirk et al., 2020). An individual’s memory capacity was defined as the largest of the two *K* measures.

### 2.5 Eye-Tracking

Eye movements were monitored using a desktop-mounted Eyelink 1000+ infrared-based eye-tracker (www.sr-research.com) with 1000 Hz sampling rate (monocular). Pupil detection was set to centroid fitting of the dominant eye. The eye-tracker was (re-)calibrated using a nine-point calibration grid at default locations. Recalibration was automatically triggered whenever a stable fixation on the fixation symbol could not be established. Trials were aborted and repeated at the end of the experiment whenever participants blinked, or their gaze was outside of a 2.5° radius around the fixation symbol, or they failed to fixate on the cued saccade target within 1000 ms after saccade cue onset.

After recording, eye tracking and EEG data were synchronized and integrated using the EYE-EEG plugin (Dimigen, Sommer, Hohlfeld, Jacobs, & Kliegl, 2011) for EEGLAB. Saccadic response times relative to the onset of saccade cues were defined as the latency of the absolute maximum of the change in horizontal gaze position (i.e. its first derivative) during the saccade interval. Microsaccades were detected using the algorithm described in Engbert and Mergenthaler, 2006: horizontal and vertical eye tracking data were smoothed to suppress noise, and eye movements were classified as saccades if they exceeded a velocity threshold *λ* of 4 standard deviations for a minimum of four samples. If multiple saccade events were detected within 100 ms, only the largest saccade from this interval was kept. Microsaccades were defined as saccades smaller than 1°.

### 2.6 EEG acquisition and preprocessing

EEG was recorded with a Biosemi Active Two EEG system (www.biosemi.nl) at 1024 Hz sampling rate from 64 Ag/AgCl electrodes arranged in a custom-made, equidistant montage, which extended to more inferior areas over the occipital lobe than the conventional 10-20 system. An additional electrode was placed below the left eye. EEG data were preprocessed using Matlab R2022b (www.mathworks.com) and the EEGLAB toolbox (Delorme & Makeig, 2004) version 2019-1.

Raw data were high-pass filtered at 0.1 Hz with a second-order Butterworth filter, low-pass filtered at 30 Hz with a Blackman FIR filter with a transition bandwidth of 5 Hz, segmented into epochs from 1000 ms before target onset until 500 ms after probe onset (5500 ms in total), and downsampled to 256 Hz. Noisy signals were defined as data of a single channel and epoch with a standard deviation larger than 2. Trials with more than 5 noisy channels were removed. If more than one quarter of all remaining trials were noisy for a given channel, that channel was interpolated for all trials using spherical-spline interpolation. Channels with fewer noisy trials were selectively interpolated on a trial-by-trial basis. EEG channels were then converted to an average reference. Epochs with signal amplitudes more extreme than *±* 400 *μ*V were rejected.

Smaller artifacts were corrected using Independent Component Analysis (extended infomax ICA). To improve the ICA solution, the original raw data were additionally high-pass filtered at 1 Hz before ICA. The resulting independent component (IC) weights were then projected back onto the original (i.e. 0.1 Hz-filtered) data. This procedure has been demonstrated to optimize detection and removal of artifactual components (Dimigen, 2020). ICs classified by the IClabel algorithm (Pion-Tonachini, Kreutz-Delgado, & Makeig, 2019) as ocular, cardiac, muscle noise, channel noise, or line noise with a probability higher than 50% were removed from the data. Finally, any remaining epochs with signal amplitudes more extreme than *±* 200 *μ*V were rejected.

To inspect the temporal and spectral distributions of the signal, the preprocessed data were transformed into the time-frequency domain by convolving the signal with a set of Morlet wavelets, defined as complex sine waves tapered by a Gaussian. The frequencies of the wavelets ranged from 2 Hz to 30 Hz. The temporal width of the wavelets was defined by the full-width at half-maximum (FWHM) for each frequency *f* as *FWHM* = 2*/f* (Cohen, 2019). To determine electrodes and frequencies of interest, lateralization of spectral power (ipsilateral - contralateral) relative to the direction of the memory-cue for left and right memory conditions was computed on no-saccade trials at all channels. For wavelet power, lateralization was computed as the difference in power at three adjacent posterior channels (white electrode markers in Figure 2C and F) in the ipsilateral minus contralateral hemisphere relative to the side indicated by the memory-cue or by the saccade-cue, respectively. Conditions without a leftward or rightward cue (i.e. target objects on the midline in experiment 1 and the no-memory control condition in experiment 2) were excluded from this computation. Since lateralization effects were specific to the alpha-band (Figure 2C and F), the single-trial EEG data were band-pass filtered from 7–13 Hz (Hamming-windowed FIR filter) and subsequently Hilbert-transformed to extract band-power for statistical analyses.

**Figure 2:**
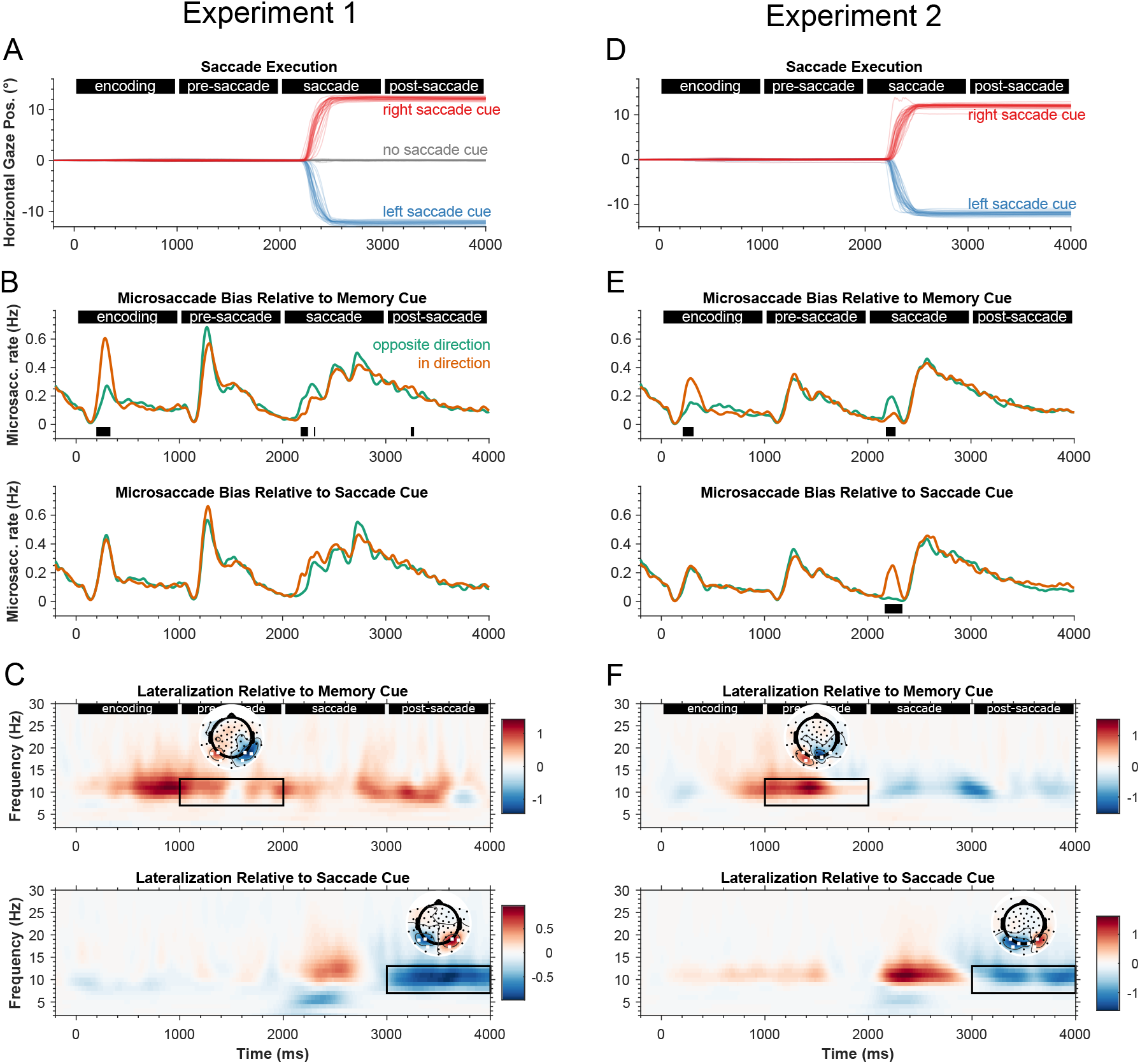
A–C: experiment 1. **A**: horizontal gaze position for each saccade-cue condition. Traces represent single subjects. **B**: directional bias of microsaccades(saccade amplitue <1°) depending on direction of memory-cue (top) and saccade cue (bottom). Note that saccades larger than 1° were not included in this analysis. **C**: top figure shows lateralization of spectral power (ipsilateral - contralateral) relative to the direction of the memory cue for left and right memory conditions on no-saccade trials. The topography shows the left-right memory difference averaged across 7–13 Hz during the pre-saccade interval. Bottom figure shows lateralization relative to the direction of the saccade cue regardless of memory condition. The topography shows the left-right saccade difference averaged across 7–13 Hz during the post-saccade interval. **D–F**: **experiment 2**, conventions as in A–C except **F** where top figure shows memory-related lateralization regardless of saccade condition.

For both the broad-band event-related potential and band-pass filtered power, lateralization was assessed with a hemispheric difference metric by subtracting the signal at left channels from the signal at a right channels (see white electrode markers in Figures 3 and 4). Note that unlike the more conventional ipsilateral-contralateral difference, it is possible to compute this right-left hemispheric difference even for conditions without a directional cue (i.e. target objects on the midline in experiment 1 and no-memory control in experiment 2). This hemispheric difference is indicative of conventional cue-related lateralization (contralateral delay activity and alpha-band lateralization) if it shows opposite polarity for leftward and rightward cues. For statistical analyses, hemispheric differences were further averaged across the pre-saccade delay interval (1000–2000 ms) and the early (3000–3500 ms) and late (3500–4000 ms) post-saccade interval.

**Figure 3:**
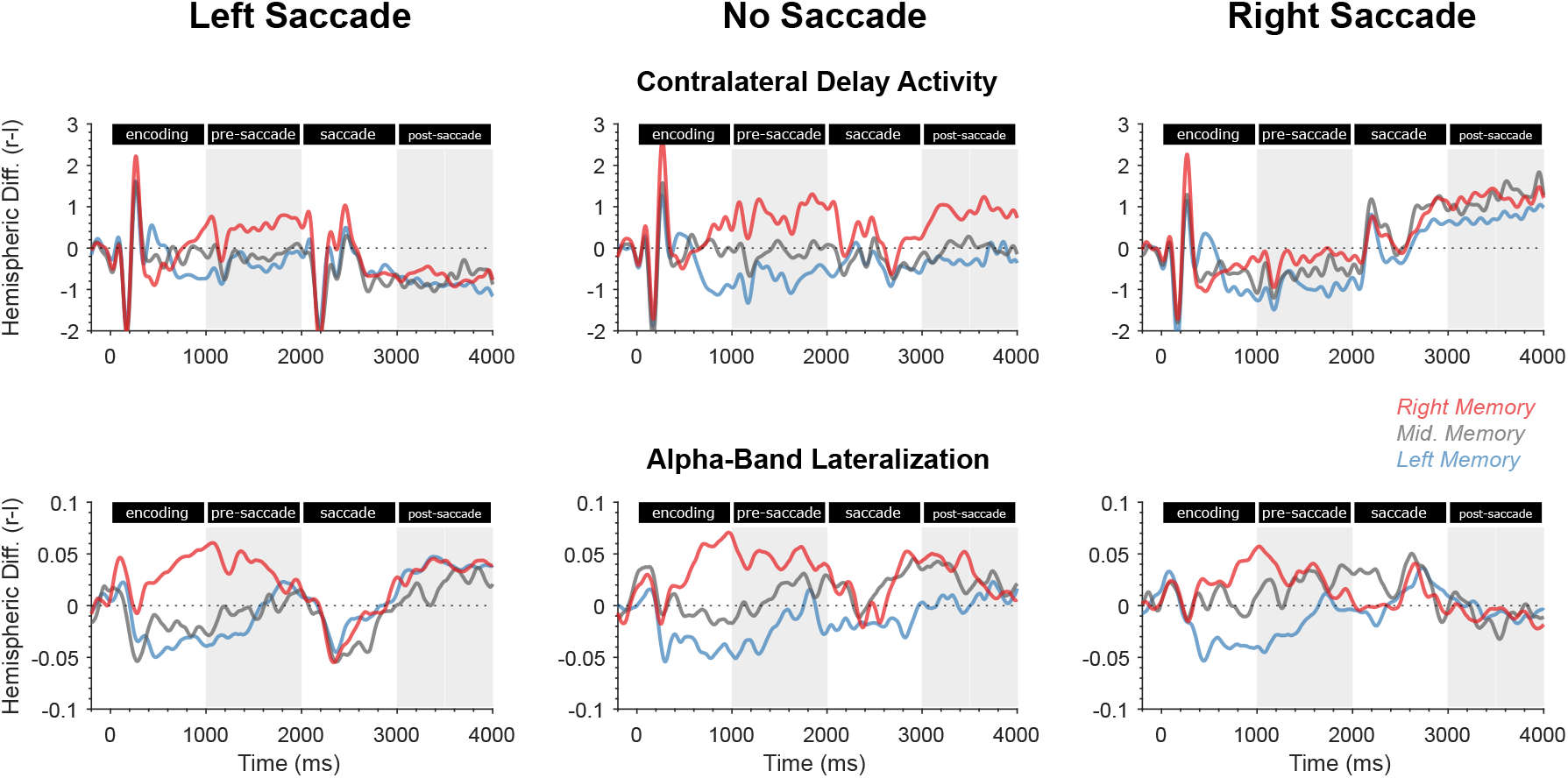
Experiment 1: hemispheric difference in event-related potentials (top) and alpha-band power (bottom). Note that this metric represents conventional contralateral delay activity and alpha-band lateralization when it is modulated by memory-cue direction.

**Figure 4:**
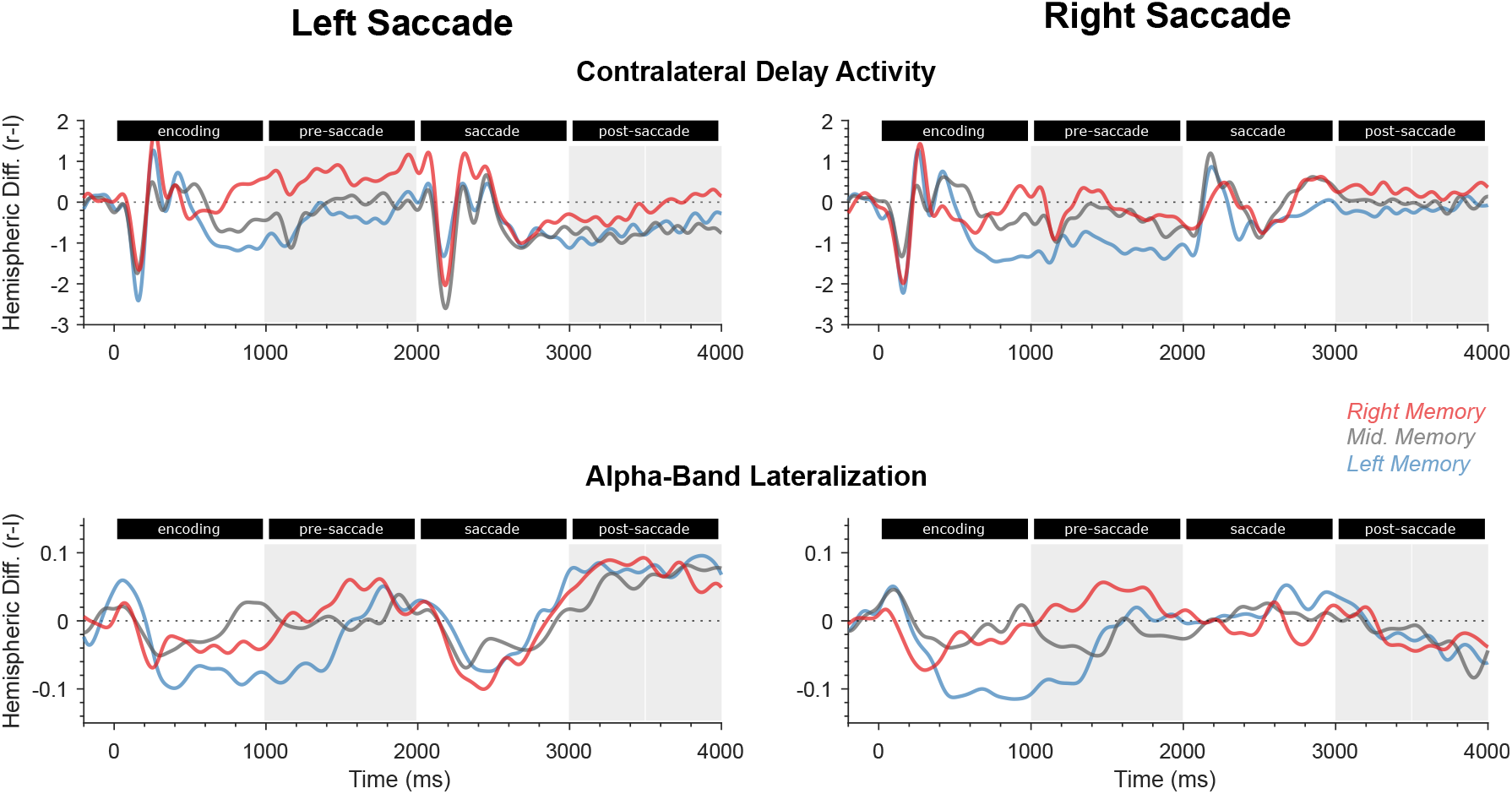
Experiment 2: hemispheric difference in event-related potentials (top) and alpha-band power (bottom). Note that this metric represents conventional contralateral delay activity and alpha-band lateralization when it is modulated by memory-cue direction.

The saccade interval was not used for statistical analysis for two reasons. First, a saccade cue appeared at the start of the saccade interval, evoking a cue-related neuronal response that would confound any saccade-related effects on the memory representation. Second, saccade execution is expected to evoke a number of ocular artifacts around the time of saccade onset. While some of these artifacts can be substantially reduced with ICA correction (Jung et al., 2000), even corrected EEG data may contain minor residual artifact traces. While such residuals are often negligible when eye movements occur infrequently and with variable timing, our design dictated that a saccade occurred on the majority of trials and with highly consistent timing across trials and subjects (See Figure 2). Therefore, even minor artifact residuals are expected to accumulate across trials. An alternative approach is to use statistical modeling to separate saccade-related signals from other signals of interest (Dandekar, Privitera, Carney, & Klein, 2012; Ehinger & Dimigen, 2019). However, this approach requires that the timing of these signals be uncorrelated across trials. As already mentioned, this was not the case here. Thus, we decided to exclude the saccade interval from statistical analysis altogether and focus only on the pre-saccade and post-saccade intervals, which were free of any (macro-)saccades.

### 2.7 Statistical Analysis

Behavioral accuracy, saccadic response times, and EEG hemispheric differences in each of the three time windows (1000–2000 ms, 3000–3500 ms, and 3500–4000 ms) were analyzed with a repeated measures analysis of variance (ANOVA) as implemented in the afex package version 1.0.1 (Singmann, Bolker, Westfall, Aust, & Ben-Shachar, 2021) for R (R Core Team, 2022). Data from experiment 1 were analyzed with a 3×3 ANOVA with factors *memory cue* (left, midline, right) and *saccade cue* (left, no-saccade, right). Data from experiment 2 were analyzed with a 3×2 ANOVA with factors *memory cue* (left, no-memory control, right) and *saccade cue* (left, right). Where appropriate, degrees of freedom and p-values were Huynh–Feldt-corrected for violations of the sphericity assumption. Pair-wise post-hoc contrasts were computed using the emmeans package version 1.6.3 for R (Lenth, 2021).

In experiment 2, the relationship between subjects’ memory capacity and lateralization in each of the three time windows was analyzed with Spearman correlations. To this end, lateralization was defined as the signal difference at ipsilateral vs. contralateral channels relative to the memory-cue (excluding the no-memory control condition) regardless of saccade-cue. Correlation coefficients were compared between the CDA and alpha-band lateralization with Dunn and Clark’s (1969) test for equality of dependent correlation coefficients (two-tailed) implemented in the cocor package version 1.1.4 for R (Diedenhofen & Musch, 2015). Since alpha-band lateralization showed a strong effect of saccade direction in the post-saccade interval, additional correlations were tested in this interval using the signal difference at ipsilateral vs. contralateral channels relative to the saccade-cue regardless of memory condition.

Directional bias of microsaccades was analyzed by comparing the rates of microsaccades in the cued direction to that of microsaccades in the opposite direction with sample-by-sample t-tests with false discovery rate correction.

## 3 Experiment 1

### 3.1 Results

#### 3.1.1 Behavior

The location of the color change was correctly located on 83% of all trials, indicating that the staircase successfully adjusted the color change magnitude. Accuracy was neither affected by the location of the memorized object (*F*(2, 58) = 1.96, *p* = .151, 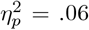), nor by saccade direction (*F*(1.49, 43.13) = 1.74, *p* = .194, 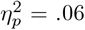), nor by their interaction (*F*(4, 116) = 0.48, *p* = .747, 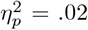). This level of accuracy required an average change of 59° in color space. The required change magnitude was neither affected by the location of the memorized object (*F*(1.00, 29.13) = 0.46, *p* = .505, 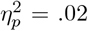), nor by saccade direction (*F*(1.00, 29.14) = 1.07, *p* = .310, 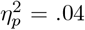), nor by their interaction (*F*(1.81, 52.53) = 1.05, *p* = .351, 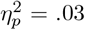).

Overall, all subjects initiated saccades promptly after saccade cue onset and refixated on the new saccade target well before the post-saccade interval started (Figure 2). For trials requiring a saccade, the average latency from saccade cue to saccade onset was 298 ms. A significant effect of memorized location (*F*(2, 58) = 13.07, *p* < .001, 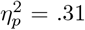) indicated that saccadic response times were fastest when a right object (293 ms) was memorized compared to left objects (297 ms) or to central objects (303 ms). There was no main effect of saccade direction (*F*(1, 29) = 1.50, *p* = .230, 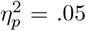). A significant interaction (*F*(2, 58) = 7.86, *p* < .001, 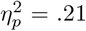) indicated that saccades were fastest when they were directed away from the memorized object, i.e. left saccades were fastest when a right object was memorized, whereas right saccades were fastest when a left object was memorized.

#### 3.1.2 CDA

##### Pre-saccade interval

During the pre-saccade delay interval, ERPs over posterior channels showed a sustained hemispheric difference whose polarity indicated the location of the cued target object. Specifically, on trials with a lateralized target, ERP amplitudes in the pre-saccade delay interval were more negative-going at contralateral channels, reflecting a conventional CDA effect. Accordingly, the hemispheric ERP amplitude difference for right channels minus left channels was more positive when a right object was memorized than when a left object was memorized (Figure 3, top row). This effect was confirmed by a significant effect of memorized location (*F*(2, 58) = 13.77, *p* < .001, 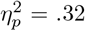). Moreover, the hemispheric difference was more positive before saccades to the left compared to the right (*F*(2, 58) = 7.41, *p* = .001, 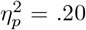) independent of memorized location. There was no interaction between memorized location and saccade direction (*F*(4, 116) = 1.12, *p* = .353, 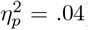).

##### Post-saccade interval

In the early post-saccade delay interval (3000 – 3500 ms), we found a similar, albeit smaller, CDA effect with the same polarity as pre-saccade, i.e. the hemispheric difference was more positive when a right object was memorized (*F*(2, 58) = 4.40, *p* = .017, 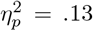). Compared to the pre-saccade interval, the effect of saccade direction reversed polarity, being more negative after saccades to the left compared to the right (*F*(2, 58) = 9.97, *p* < .001, 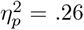). There was no interaction between memorized location and saccade direction (*F*(3.20, 92.92) = 0.63, *p* = .609, 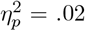).

In the late post-saccade interval (3500 – 4000 ms), the main effect of memorized location was no longer significant (*F*(2, 58) = 2.28, *p* = .111, 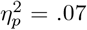), whereas the effect of saccade direction still was (*F*(2, 58) = 16.78, *p* < .001, 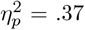). A post-hoc test revealed a weak effect of memorized location for trials without a saccade (*t*(29) = *−*2.095, *p* = .045). Again, there was no interaction between memorized location and saccade direction (*F*(2.51, 72.75) = 1.09, *p* = .353, 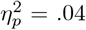).

#### 3.1.3 Alpha lateralization

A wavelet-based time-frequency analysis confirmed that for lateralized targets, alpha power was reduced in the hemisphere contralateral to the memorized object relative to the ipsilateral hemisphere during encoding and during the pre-saccade delay interval, especially on trials without saccade. This effect was most pronounced at frequencies from 7 to 13 Hz and at posterior channels (Figure 2).

##### Pre-saccade interval

During the pre-saccade delay interval, posterior alpha-band power showed a sustained hemispheric difference depending on the location of the memorized target object. Specifically, on trials with a lateralized target, alpha-band power was reduced at contralateral compared to ipsilateral channels, reflecting a conventional memory-related lateralization effect. Accordingly, the hemispheric difference for right channels minus left channels was more positive when a right object was memorized than when a left object was memorized (Figure 3, bottom row).

This effect was confirmed by a significant effect of memorized location on the hemispheric difference waveform (*F*(2, 58) = 23.21, *p* < .001, 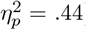). The hemispheric difference waveform was not affected by the direction of the upcoming saccade (*F*(2, 58) = 0.18, *p* = .836, 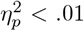) and there was no interaction between memorized location and saccade direction (*F*(4, 116) = 2.30, *p* = .063, 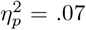).

##### Post-saccade interval

In the early post-saccade delay interval (3000 – 3500 ms), there was no main effect of memorized location on alpha-band lateralization (*F*(2, 58) = 1.29, *p* = .284, 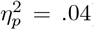). However, an interaction between memorized location and saccade direction (*F*(3.31, 96.13) = 3.21, *p* = .023, 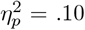) indicated an effect of memorized location on trials without saccade (*t*(29) = *−*2.920, *p* = .179), but not for trials with a left (*t*(29) = *−*2.920, *p* = .979) or a right-going saccade (*t*(29) = 0.537, *p* = .853) Moreover, lateralization was affected by saccade direction such that alpha-band power was stronger at channels ipsilateral to saccade direction (*F*(2, 58) = 4.00, *p* = .024, 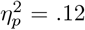).

In the late post-saccade delay interval (3500 – 4000 ms), we found neither a main effect of memorized location (*F*(1.76, 50.95) = 0.09, *p* = .894, 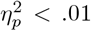) nor an interaction effect (*F*(4, 116) = 0.34, *p* = .852, 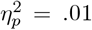), but a sustained effect of saccade direction (*F*(1.64, 47.53) = 5.44, *p* = .011, 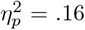).

#### 3.1.4 Microsaccade rates

The direction of microsaccades was significantly biased in the direction of the memory-cue during the early encoding interval from 195 to 332 ms, and in the opposite direction during the saccade interval from 2175 to 2242 ms (Figure 2B, top). In this interval, most microsaccades were small corrective saccades that occurred soon after the eyes had landed on the saccade target. During the post-saccade interval, microsaccades were briefly biased in the direction opposite to the memory-cue from 3242 to 3273 ms. Microsaccade rates were not significantly biased by the saccade-cue (Figure 2B, bottom) in any time window.

### 3.2 Discussion

Memory accuracy was not affected by the location of the memorized object nor by saccade direction, implying that EEG effects between memory or saccade conditions were not confounded by differences in task difficulty.

During the encoding and pre-saccade intervals, ERP amplitudes were more negative over the hemisphere contralateral to the location indicated by the memory cue than over the ipsilateral hemisphere. Accordingly, the hemispheric difference metric (right - left channels) was more negative-going for left memory-cues, more positive-going for right memory-cues, and near zero for midline cues (Figure 3). Thus, for lateralized targets, our result are equivalent to the more conventional difference between contralateral and ipsilateral channels reported in previous studies on the CDA (Luria et al., 2016; Vogel & Machizawa, 2004). In the post-saccade interval, the magnitude of the CDA was overall reduced. A similar decay has been noted in many previous CDA studies (Roy & Faubert, 2023) even across much shorter delay intervals than the three-second delay in the present study. While some of this reduction could be attributed to degradation of the memory representation over time, good behavioral performance in all experimental conditions implies that memories were indeed maintained in the majority of trials. Moreover, on trials without a saccade, a substantial CDA was observed until the very end of the delay interval, implying that the CDA is not generally too short-lived. While the amplitude reduction was stronger after a saccade, the crucial finding is that the *polarity* of the post-saccade CDA was identical to that of the pre-saccade CDA regardless of saccade direction. In other words, the involvement of the two hemispheres in maintaining the memory representation was still determined by the target object’s original retinotopic location at encoding, not by its updated spatiotopic location after the saccade. Importantly, the probe stimulus used to test subjects’ memory performance was not lateralized, but centered on the new fixation (see probe in Figure 1), rendering it unlikely that post-saccade lateralization merely reflects anticipation of an upcoming lateralized stimulus. Rather, this finding indicates that the CDA reflects a memory representation coded in a retinotopic frame of reference (Golomb & Kanwisher, 2012b; Shafer-Skelton & Golomb, 2018).

The CDA also showed a strong effect of saccade direction (Figure 3). This was most notable in the post-saccade interval where a slow, direct current offset was superimposed on the memory-related CDA effect. This reflects the well-known corneo-retinal artifact, whereby a saccade and subsequent sustained fixation on a lateral saccade target leads to a sustained shift of the corneo-retinal electric potential (Dandekar et al., 2012; Lins, Picton, Berg, & Scherg, 1993). Curiously, the results showed an effect of saccade direction already in the pre-saccade interval. Since the direction given by the saccade-cue was unpredictable, this likely represents a signal-processing artifact due to high-pass filtering. Note that the polarity of the pre-saccade effect is opposite of that of the post-saccade corneo-retinal artifact and that the polarity reversal occurs with the sharp transient signal at the start of the saccade-interval (approximately 2200 ms). Similar filtering artifacts under such conditions have been reported by Tanner, Morgan-Short, and Luck, 2015 and Acunzo, MacKenzie, and van Rossum, 2012. Importantly, there was no interaction between the saccade-related artifact and the memory-related CDA effect, ruling out this signal-processing artifact as a confound.

Alpha-band power in the pre-saccade interval showed a conventional topographic pattern that was lateralized in the direction of the memory-cue. Specifically, power at frequencies around 10 Hz was reduced at channels over the hemisphere contralateral to the location indicated by the memory-cue (Figure 2C, top panel). Accordingly, the hemispheric difference metric was more negative going for left memory cues, more positive going for right memory cues, and near zero for midline cues (Figure 3). This pattern paralleled the CDA effect in the pre-saccade interval and is generally consistent with previous findings (Fukuda, Kang, & Woodman, 2016; Sauseng et al., 2009; Schroeder, Aagten-Murphy, & Busch, 2023) showing that the polarity of alpha-band lateralization tracks the attentional prioritization of memory-relevant external locations and internal stimulus representations (van Ede, 2018). In no-saccade trials, this conventional lateralization towards the memorized object continued throughout the delay interval up to the onset of the probe stimulus. In saccade trials, however, post-saccade lateralization reflected only the direction of the saccade (Figure 2C, bottom panel). For example, after a saccade to the left, alpha-band power was reduced over the left hemisphere compared to the right hemisphere, consistent with prioritization of locations near the initial fixation during encoding.

It is important to stress that during the post-saccade interval, there were no stimuli near the initial central fixation marker that might have attracted subjects’ attention. Moreover, subjects were required to keep fixating on the new lateral fixation marker and wait for the appearance of the probe stimulus, which was a single object centered on the new fixation marker. Given that subjects had to focus attention at the new fixation for performing the color change detection task, it is very unlikely that the lateralization pattern in the post-saccade interval reflects a covert shift of attention in anticipation of an upcoming saccade back to the initial fixation.

Could post-saccade lateralization, instead, reflect involuntary eye movements? Although we monitored eye movements to make sure that fixations and cued saccades were executed as instructed (Figure 2A), we could not rule out that subjects made occasional, possibly involuntary miniature saccades not exceeding our criterion for trial exclusion. The direction of such microsaccades is often biased in the direction of covertly attended locations (Hafed & Clark, 2002; Rolfs, 2009) or of memorized objects (van Ede, Chekroud, & Nobre, 2019); that is, in the same general direction as alpha-band lateralization. Moreover, trial-by-trial variations in the amplitude and timing of alpha lateralization are tightly coupled with microsaccades (Liu, Nobre, & van Ede, 2022, 2023a). In this experiment, microsaccades were indeed biased in the direction of the cued object during the early encoding interval (app. 200–300 ms), coinciding approximately with the onset of alpha-band lateralization in the same direction (Figure 2B, top and 2C, top). However, while alpha power remained lateralized throughout the encoding and pre-saccade delay interval, no further microsaccade bias was observed at longer latencies. During the post-saccade interval, while alpha power was strongly lateralized towards the memorized object on no-saccade trials and towards the initial fixation on saccade trials, microsaccades did not show any systematic memory-related or saccade-related directional bias (Figure 2B, bottom and 2C, bottom).

These findings could suggest that the transition from lateralization directed towards the memorized object before the saccade to a lateralization towards the initial fixation after the saccade might reflect spatiotopic updating, whereby memory-related attentional prioritization is “bound” to the memorandum’s real world location during encoding (Brincat et al., 2021). Alternatively, the post-saccade lateralization towards the initial, central fixation location might reflect a memory-unspecific bias towards previously relevant or currently salient locations. This interpretation would predict post-saccade lateralization towards the initial fixation marker even if the task does not require any memory maintenance at all. This prediction was tested in experiment 2, which included a no-memory control condition.

## 4 Experiment 2

Experiment 2 was conducted to replicate findings of the first experiment, and to test whether post-saccade alpha-band lateralization opposite to saccade direction, i.e. towards the initial fixation location, was related to maintenance of memorized objects. Experiment 2 was largely similar to the first experiment (Figure 1B), but included an additional control condition: in the no-memory condition, all objects were uniformly white and subjects were not required to remember any of them. Accordingly, the probe stimulus was not used for a color memory task, but for a perceptual color discrimination task.

To further probe the mnemonic relevance of CDA and alpha-band lateralization, we correlated these lateralized EEG indices with behavioral short-term memory capacity. The design of the main task was not suited for estimating memory capacity for several reasons. First, capacity estimates are only valid if set size is as large as or larger than the true capacity (Rouder et al., 2011). Thus, a set size of two (experiment 1) or three (experiment 2) may underestimate capacity for some subjects. Second, capacity is estimated from accuracy scores. However, the main task included a staircase procedure that adjusted task difficulty in order to keep accuracy at a fixed level for all subjects and conditions. Finally, capacity estimates based on change detection tasks are only valid if target and foil colors are highly distinguishable (Rouder et al., 2011). However, due to the staircase procedure the target-foil color difference was variable across trials and included less distinguishable colors. Therefore, experiment 2 included an additional assessment of memory capacity using a procedure adapted from Brady et al., 2016 and Quirk et al., 2020.

### 4.1 Results

#### 4.1.1 Behavior

Color changes were correctly localized for 76% of left memorized objects, and for 75% of right memorized objects. In the no-memory control condition, the odd color in the probe object was correctly localized on 87% of all trials. The ANOVA yielded a main effect of memory condition (*F*(1.21, 41.27) = 18.49, *p* < .001, 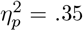). Contrasts showed that this effect was due to better accuracy in the no-memory control condition compared to left memorized objects (*p* = .006) and compared to right memorized objects (*p* = .004). Accuracy was not affected by saccade direction (*F*(1, 34) = 1.35, *p* = .254, 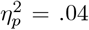), nor by an interaction between memory and saccade direction (*F*(1.74, 59.03) = 1.62, *p* = .209, 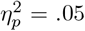).

This level of accuracy required an average change in color space of 136° for left memorized objects, 135° for right memorized objects, and 25° in the no-memory control condition. The ANOVA yielded a main effect of memory condition (*F*(1.00, 34.07) = 104.64, *p* < .001, 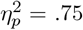). Contrasts showed that this effect was due to smaller color change in the no-memory control condition compared to left memorized objects (*p* < .001) and compared to right memorized objects (*p* < .001). Color change was not affected by saccade direction *F*(1, 34) = 0.01, *p* = .908, 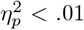), nor by an interaction between memory and saccade direction (*F*(1.01, 34.26) = 1.29, *p* = .265, 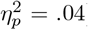).

Saccadic response times were slightly faster for left saccades (281 ms) than for right saccades (287 ms), as indicated by a significant effect of saccade direction (*F*(1, 34) = 4.57, *p* = .040, 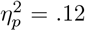). There was no effect of memory condition (*F*(1.65, 56.17) = 1.25, *p* = .289, 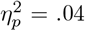), but a significant interaction between memory and saccade direction (*F*(1.49, 50.58) = 4.23, *p* = .030, 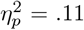), indicating that saccades were fastest when they were directed away from a memorized object.

Individual memory capacity *K* ranged from 1.8 to 5.3 (mean 3.3).

#### 4.1.2 CDA

##### Pre-saccade interval

Similar to experiment 1, ERPs showed a sustained hemispheric difference whose polarity indicated the location of the cued target object, reflecting a conventional CDA effect (Figure 4, top row). This was confirmed statistically by a significant effect of memorized location (*F*(2, 68) = 15.70, *p* < .001, 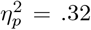). Moreover, the hemispheric difference was more positive on trials with saccades to the left compared to the right (*F*(1, 34) = 16.22, *p* < .001, 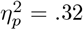). There was no interaction between memorized location and saccade direction (*F*(2, 68) = 2.41, *p* = .098, 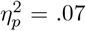). Subjects with stronger lateralization, i.e. more negative amplitudes over the hemisphere contralateral to the memorized object, had higher memory capacity (Spearman’s *ρ* = -.52, *p* = .001).

##### Post-saccade interval

In the early post-saccade delay interval (3000 – 3500 ms), we found a similar, albeit smaller, CDA effect with the same polarity as pre-saccade (*F*(2, 68) = 4.81, *p* = .011, 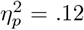). The hemispheric difference was more negative on trials with saccades to the left compared to the right (*F*(1, 34) = 17.59, *p* < .001, 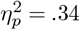). There was no interaction between memorized location and saccade direction (*F*(2, 68) = 0.36, *p* = .696, 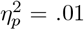). Subjects with stronger lateralization in the early post-saccade interval had higher memory capacity (Spearman’s *ρ* = -.45, *p* = .007).

In the late post-saccade interval (3500 – 4000 ms), we found sustained main effects of memorized location (*F*(1.69, 57.34) = 4.66, *p* = .018, 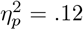) and saccade direction (*F*(1, 34) = 4.33, *p* = .045, 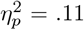) There was no interaction between memorized location and saccade direction (*F*(2, 68) = 1.20, *p* = .308, 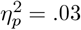). Subjects with stronger lateralization in the late post-saccade interval had higher memory capacity (Spearman’s *ρ* = -.41, *p* = .015).

#### 4.1.3 Alpha lateralization

##### Pre-saccade interval

Similar to experiment 1, alpha-band power during the pre-saccade delay interval showed a sustained hemispheric difference depending on the location of the memorized target object (Figure 4, bottom row; *F*(2, 68) = 7.09, *p* = .002, 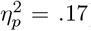). The hemispheric difference was not affected by the direction of the upcoming saccade (*F*(1, 34) = 1.85, *p* = .182, 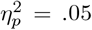) and there was no interaction between memorized location and saccade direction (*F*(2, 68) = 0.88, *p* = .419, 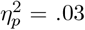). The magnitude of memory-related lateralization was not correlated with memory capacity (Spearman’s *ρ* = -.08, *p* = .658) and was significantly smaller than that between CDA and capacity (*p* = .047).

##### Post-saccade interval

In the early post-saccade delay interval (3000 – 3500 ms), there was no main effect of memorized location on alpha-band lateralization (*F*(2, 68) = 1.01, *p* = .371, 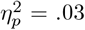), but a strong effect of saccade direction (*F*(1, 34) = 17.07, *p* < .001, 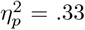). There was no significant interaction between memorized location and saccade direction (*F*(2, 68) = 0.77, *p* = .466, 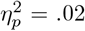). Contrasts showed that the difference between left and right going saccades was significant for left memorized objects (*t*(34) = 3.107, *p* = .003), for right memorized objects (*t*(34) = 3.374, *p* = .002), and even for the no-memory control condition (*t*(34) = 3.374, *p* = .025). The magnitude of memory-related lateralization was not correlated with memory capacity (Spearman’s *ρ* = .11, *p* = .535) and was significantly smaller than that between CDA and capacity (*p* = .030). Likewise, saccade-related alpha-band lateralization was not correlated with capacity (Spearman’s *ρ* = -.10, *p* = .575).

In the late post-saccade delay interval (3500 – 4000 ms), we found neither a main effect of memorized location (*F*(2, 68) = 0.06, *p* = .941, 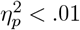) nor an interaction effect (*F*(2, 68) = 0.42, *p* = .658, 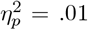), but a sustained effect of saccade direction (*F*(1, 34) = 41.42, *p* < .001, 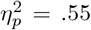). Contrasts showed that the difference between left and right going saccades was significant for left memorized objects (*t*(34) = 5.382, *p <* .001), for right memorized objects (*t*(34) = 5.239, *p <* .001), and even for the no-memory control condition (*t*(34) = 4.038, *p <* .001). The magnitude of memory-related alpha-band lateralization was not correlated with memory capacity (Spearman’s *ρ* = -.02, *p* > .915), but this correlation was not significantly smaller than that of the CDA in this interval (*p* = .117). Likewise, saccade-related alpha-band lateralization was not correlated with capacity (Spearman’s *ρ* = -.10, *p* = .569).

#### 4.1.4 Microsaccade rates

The direction of microsaccades was significantly biased in the direction of the memory-cue during the early encoding interval from 207 to 312 ms, and in the opposite direction during the saccade interval from 2175 to 2269 ms (Figure 4E, top). As in experiment 1, most microsaccades in this interval were small corrective saccades that occurred soon after the eyes had landed on the saccade target. No memory-related bias occurred in the post-saccade interval. Microsaccades were additionally biased in the direction of the saccade cue from 2164 to 2335 ms (Figure 4E, bottom).

### 4.2 Discussion

Experiment 2 largely replicated the findings of the first experiment. In the pre-saccade interval, CDA and alpha-band power showed the conventional lateralization towards the memorized object. In the post-saccade interval, the CDA maintained this direction of lateralization independent of saccade direction. By contrast, post-saccadic alpha-band lateralization was again memory-unspecific, but instead directed towards the initial fixation location. Experiment 2 extends this finding by showing this effect of saccade direction on post-saccade alpha lateralization even in the no-memory control condition. Further supporting this memory-unspecific lateralization was the finding that alpha-band lateralization was not correlated with individual short-term memory capacity, while both pre-saccade and post-saccade CDA amplitudes were strongly correlated with capacity. Together, these findings argue against a direct role of alpha-lateralization in memory-related attentional prioritization of memoranda’s original locations. Instead, they suggest that post-saccade alpha-band lateralization reflects a memory-unspecific bias towards the screen center, whereas the CDA reflects memory representations coded in a retinotopic frame of reference that is not updated after an eye movement.

## 5 General Discussion

Visual short-term memory maintains visual information while the corresponding stimulus is no longer present. Neuroimaging and single-unit studies have demonstrated that visual short-term memory storage is mediated by sensory recruitment or sustained activity in posterior cortical regions that initially encoded the memorized stimulus (Pasternak & Greenlee, 2005; Postle, 2006). Each cerebral hemisphere contains a visual short-term memory store for the respective contralateral hemifield. These lateralized stores can operate largely independently, as the number of memorized objects in one hemifield has little effect on storage capacity (Delvenne, 2005; Umemoto et al., 2010) and neural correlates of memory load (Kornblith et al., 2016) in the opposite hemisphere. Accordingly, when to-be-remembered stimuli are exclusively located within one visual hemifield, neural correlates of short-term memory maintenance are lateralized to the contralateral hemisphere. This study investigated whether lateralized electrophysiological indices of visual short-term memory maintenance – contralateral delay activity (CDA) and alpha-band lateralization – are lateralized towards the memoranda’s original location on the retina or towards their updated spatiotopic location after an eye movement during the maintenance interval. To this end, subjects encoded color stimuli in the left or right hemifield and made a lateral saccade during the memory maintenance interval. Thereby, a location that was originally on the left of fixation during encoding came to be situated to the right of fixation after a leftward saccade. After the maintenance interval ended, participants performed a color-change detection task at the new fixation (Figure 1).

### 5.1 Post-saccade CDA reflects a retinotopic reference frame

CDA conventionally reflects a more negative-going signal at posterior channels contralateral to the encoded stimulus during the maintenance interval (see top center panel in Figure 3). In both experiments, we found a conventional CDA throughout the three-seconds-long delay interval in trials without a saccade and in the pre-saccade delay interval in trials with a saccade (see top center panel in Figure 3). Importantly, in trials with a leftward or rightward saccade, the polarity of the post-saccade CDA did not change with respect to the pre-saccade CDA. Specifically, there was no polarity-inversion when a saccade brought the location where the memorized object had been presented at encoding into the opposite hemifield (see top left and right panels in Figure 3 and 4). Moreover, the magnitude of both pre-saccade and post-saccade CDA was correlated with subjects’ individual memory capacity estimated from an independent memory task. Since the post-saccade delay interval was followed by a non-lateralized probe object that was centered at the post-saccade fixation, the post-saccade CDA lateralization cannot reflect anticipation of an upcoming lateralized probe stimulus. This finding corresponds well with previous studies demonstrating that the CDA specifically reflects selective maintenance in visual short-term memory rather than general attentional orienting towards anticipated events (Hakim, Adam, Gunseli, Awh, & Vogel, 2019), even when the task does not explicitly require memorization of the targets’ spatial arrangement (Kuo, Stokes, & Nobre, 2012), and that its polarity is determined by the target’s location at encoding and not by the anticipated location of the probe (Grubert & Eimer, 2015). Our finding that the polarity of the post-saccade CDA is independent of saccade direction therefore indicates that the CDA reflects maintenance of a memory representation that is coded in retinotopic rather than spatiotopic coordinates relative to the memoranda’s location at encoding.

### 5.2 Alpha-band lateralization reflects memory-independent screen bias

Alpha-band lateralization in visual short-term memory tasks conventionally reflects relatively reduced power at posterior channels contralateral to the encoded stimulus during memory maintenance. Indeed, we found this conventional pattern in the early delay interval before saccade onset and in the late delay interval in trials without a saccade in experiment 1 (see Figure 2C and bottom center panel in Figure 3). In saccade trials, post-saccade alpha-band lateralization reflected only the direction of the saccade, being directed “against” saccade direction. This result is partially compatible with a spatiotopic updating of the target’s original location. However, while post-saccade lateralization reflected the previous location of the entire stimulus array (at the screen center) during encoding, it did not track the target’s specific hemifield within the stimulus array. Given that after, say, a left saccade, the screen locations of left and right targets would both be situated at different eccentricities within the right hemifield, it is possible that the spatial resolution of EEG is insufficient for resolving subtle differences in post-saccade lateralization, thus explaining why we failed to find an effect of target hemifield after a saccade.

An alternative interpretation of this result is that post-saccade alpha lateralization does not actually reflect a process that is relevant for memory maintenance. Note that, in keeping with conventional paradigms designed to evoke CDA and alpha-band lateralization, the target’s original locations at encoding were always to the left or right of the initial fixation at the screen center. Thus, lateralization “against” saccade direction is equivalent to lateralization “towards” the screen center. Contradicting the mnemonic relevance of this post-saccade lateralization, we found the same lateralization towards the screen center even in the no-memory control condition in experiment 2 (Figure 4), in which target locations were entirely blank during “encoding”. Moreover, neither the conventional pre-saccadic lateralization towards the memorized location nor the post-saccadic lateralization towards the screen center were correlated with memory performance across subjects. In light of this finding, it appears unlikely that post-saccade alpha-band lateralization reflects a spatiotopically updated memory trace, but a memory-independent bias towards the screen center.

Numerous studies have demonstrated alpha-band lateralization induced by task-relevant cues or events. For instance, task-relevant and informative cues indicating a to-be attended location or a to-be-memorized object induce a typical power decrease contralateral to the cued location (Foxe & Snyder, 2011). Since the alpha rhythm is indicative of a state of physiological inhibition, such lateralization has been interpreted as an active gating mechanism supporting the selective amplification of relevant and/or inhibition of irrelevant information (Jensen & Mazaheri, 2010; van Ede, 2018). However, other findings suggest that alpha lateralization is more generally driven by conspicuous or salient events, even when not task-relevant. Several studies have shown cue-induced alpha lateralization even for cues that were not temporally (Green & McDonald, 2010) or spatially (Günseli et al., 2019) informative of the relevant locations. Moreover, several studies have demonstrated that task-irrelevant distractors can induce lateralization towards those events indicating attentional capture instead of lateralization away from them indicating inhibition (Schroeder, Ball, & Busch, 2018; Schroeder et al., 2023; Hakim, Feldmann-Wüstefeld, Awh, & Vogel, 2021; Keefe & Störmer, 2021; Balestrieri, Michel, & Busch, 2022). Finally, several studies have found that the timing of alpha lateralization can deviate from the expected time course of attention orienting: it outlasts other electrophysiological and behavioral effects of attentional orienting, and it can even outlast the subject’s behavioral response (Keefe & Störmer, 2021; Antonov, Chakravarthi, & Andersen, 2020; Bacigalupo & Luck, 2018). In sum, these findings demonstrate that alpha-band lateralization can reflect a lingering trace of excitation due to salient events, even if these events are not directly task-relevant. It is therefore possible that in the present study, the screen center was such a conspicuous location, resulting in post-saccade alpha lateralization towards the screen center that was dissociated from the current focus of spatial attention at the new fixation.

Alternatively, it is possible that the saccade disrupted the mnemonic role of alpha lateralization during fixation, even though behavioral retention performance remained largely intact. For instance, Bullock et al. (2023) used inverted encoding modeling to reconstruct the location of the memorized object from the topographical distribution of alpha power. The reconstruction of memorized location was disrupted by saccades during the delay interval, even when the final saccade returned to the initial fixation location. Thus, our findings may not necessarily invalidate the functional relevance of alpha lateralization for memory maintenance in studies using continuous fixation.

Recent studies have found that the coincidence of spatial cues and alpha-band lateralization may be mediated by a cue-induced bias in the direction of miniature eye movements (Liu et al., 2022; Popov, Gips, Weisz, & Jensen, 2022; Liu, Nobre, & van Ede, 2023b). Our analysis of the eye tracking data showed that subjects executed the instructed saccades accurately within the saccade interval. Importantly, microsaccades were not systematically biased in any particular direction during the pre-saccade and post-saccade intervals (Figure 2B, E), making it unlikely that alpha-band lateralization in those intervals was a result of a directional bias of miniature eye movements.

### 5.3 Reference frames in visual short-term memory

Our paradigm resembles that of a study by Brincat et al. (2021) on interhemispheric remapping of visual short-term memories. There, macaques were trained to memorize a lateralized target object and then make a saccade bringing the target’s location into the opposite visual hemifield. In the interval before the saccade, firing rates were stronger and more informative in lateral prefrontal cortex of the contralateral hemisphere. In the interval after the saccade, this lateralization inverted, consistent with transfer of the memory trace between hemispheres and a spatiotopic reference frame coding locations in world- or screen-centered coordinates. By contrast, behavioral studies by Golomb and Kanwisher (2012b) and Shafer-Skelton and Golomb (2018) showed that humans are actually worse at reporting where a memorized target had been located on the screen than reporting where it had been located on the retina, and that spatiotopic memory deteriorated more with each intervening saccade. Likewise, studies on the reference frame of sustained spatial attention have demonstrated that attention is maintained primarily in retinotopic coordinates immediately after saccades. Specifically, Golomb et al. (2008) had subjects execute saccades while sustaining attention at another location. Behavioral performance, tested with probe stimuli after the saccade, was facilitated for probes presented at the previously attended retinotopic location, but not at the same spatiotopic location on the screen. Thus, while our paradigm and rationale is more similar to Brincat et al. (2021), our finding that the polarity of CDA lateralization reflects only the target’s retinotopic hemifield at encoding, but not its updated screen location supports the view that visual short-term memory representations are encoded and maintained in retinotopic coordinates (Golomb & Kanwisher, 2012b).

A potential reason why we did not find evidence for a transfer of memory representations between hemispheres following a saccade could be due to the different neural sources involved. While Brincat et al. (2021) recorded from monkey prefrontal cortex, the topography of the CDA is not indicative of a prefrontal source. In fact, two MEG studies have localized the cortical sources underlying the CDA to posterior parietal cortex (Robitaille, Grimault, & Jolicœur, 2009; Becke, Müller, Vellage, Schoenfeld, & Hopf, 2015). It is therefore possible that posterior perceptual areas code visual short-term memories in a retinotopic reference frame, such that sustained internally or externally directed attention can linger in retinotopic coordinates for several hundred milliseconds after the execution of a saccade, while higher level areas may be involved in updating these representations to spatiotopic coordinates that are more relevant for interacting with the external world.

## Acknowledgements

We thank Teresa Berther and Johanna Seroka for help with data acquisition, and Laura Huber, Charlotte Meinke, and Franziska Rarey for assisting with the piloting phase of the study.

## Data availability

Data will be made available upon acceptance of the manuscript under https://osf.io/x4y9n/ (Open Science Framework).

